# A translational kidney organoid system bolsters human relevance of clinical development candidate

**DOI:** 10.1101/2019.12.30.891440

**Authors:** Amy Westerling-Bui, Thomas W. Soare, Srinivasan Venkatachalan, Michael DeRan, Eva Maria Fast, Alyssa B. Fanelli, Sergii Kyrychenko, Hien Hoang, Grinal M. Corriea, Wei Zhang, Maolin Yu, Matthew Daniels, Goran Malojcic, Xin-Ru Pan-Zhou, Mark W. Ledeboer, Jean-Christophe Harmange, Maheswarareddy Emani, Thomas T. Tibbitts, John F. Reilly, Peter Mundel

## Abstract

A major challenge in drug discovery is gaining confidence in the human relevance of pre-clinical animal studies. While human iPSC-derived organoids offer exciting opportunities to address this, concerns about applicability and scalability remain. Here, we report a high-throughput organoid platform for assessment of kidney disease targeting compounds in a human system. We confirmed platform reproducibility by single cell RNA-Seq (scRNA-Seq) and derived a NanoString panel for efficient quality control (QC). Organoid transplantation in rats for 2 to 4 weeks promoted organoid maturation and vascularization. In functional studies, cyclosporine A (CsA) and GFB-887, a novel TRPC5 channel blocker, protected kidney organoids from injury. Pharmacodynamic studies with GFB-887 delivered orally to rats were also successfully performed in human transplanted organoids. These data show how human organoids can deliver confidence in taking development candidate compounds to the clinic, fulfilling their promise to revolutionize drug discovery.

## Introduction

Despite exciting biopharmaceutical advances against significant unmet medical needs, there remains a substantial imperative to discover new treatments for intractable diseases, including chronic kidney disease (CKD). One in nine people worldwide suffer from CKD making it a global epidemic affecting more than 850 million people (Jager et al., 2019). CKD represents many disorders, each with diverse, multifactorial genetic and environmental drivers. As a consequence, drug development in the kidney space has been hampered by limited translatability of animal models to the human condition (Inrig et al., 2014).

New opportunities for human preclinical target evaluation have arisen through the development of iPSC-derived organoids including kidney organoids grown *in vitro* (Dvela-Levitt et al., 2019; Morizane et al., 2015; Taguchi and Nishinakamura, 2017; Takasato et al., 2016b) or transplanted *in vivo* (Sharmin et al., 2016; Subramanian et al., 2019; van den Berg et al., 2018). However, the use of organoids in drug discovery has been limited by problems with scalability, and by the fact that organoids grown *in vitro* could not be used for pharmacokinetic (PK) and pharmacodynamic (PD) studies. Here we describe a scalable high-throughput organoid platform for human preclinical assessment of kidney disease-targeting drugs. We demonstrate the utility of the platform in three examples: (i) protection of the podocyte cytoskeleton by the calcineurin inhibitor CsA, which is clinically used to treat kidney diseases (Mathieson, 2008); (ii) preclinical evaluation of GFB-887, a novel small molecule TRPC5 ion channel blocker in kidney organoids *in vitro*, and (iii) PK/PD studies with GFB-887 in organoids transplanted in rats. For the pharmacodynamic studies, we chose the protamine sulfate (PS) model of podocyte injury because (i) it has been widely used in rats (Kerjaschki D, 1978; Seiler et al., 1975), (ii) PS has been shown to activate TRPC5 (Schaldecker et al., 2013) and (iii) TRPC5 inhibition has been previously shown to be protective using this model (Schaldecker et al., 2013). Organoid reproducibility and differentiation were monitored by scRNA-Seq analysis and these results informed the design of a NanoString panel for rapid and efficient QC. Our studies illuminate how organoids can be harnessed for human preclinical target validation and compound evaluation.

## Results

### Development of a high-throughput *in vitro* kidney organoid platform

Building on prior seminal work (Morizane and Bonventre, 2017; Takasato et al., 2016a; Takasato et al., 2016b), we established a high-throughput system for human kidney organoids, by developing a scalable, modified version of previously described protocols (Takasato et al., 2016a; Takasato et al., 2016b) (Fig. 1A). The modifications, which included the seeding of iPSCs at 275,000 cells per T25 flask and treating them with 10μM and 8 μM CHIR99021 on day 0 and day 2, respectively, enabled the reproducible, high-throughput (> 500 organoids/week) generation of human kidney organoids, thereby making their use suitable for drug discovery (Fig. 1A). In keeping with published results (Morizane et al., 2015; Subramanian et al., 2019; Takasato et al., 2016b), the organoids expressed protein markers of podocytes (synaptopodin), proximal tubule (LTL), and distal (E-cadherin) tubules (Fig. 1A, B). Of note, *in vitro* D28 organoids also contained endothelial marker (CD31/PECAM1) expressing cells that formed vessel structures. In addition, most of the nephrons were concentrated in one plane at a depth of around 150μm from the apical side of the organoid (Fig. 1C).

**Figure 1.**
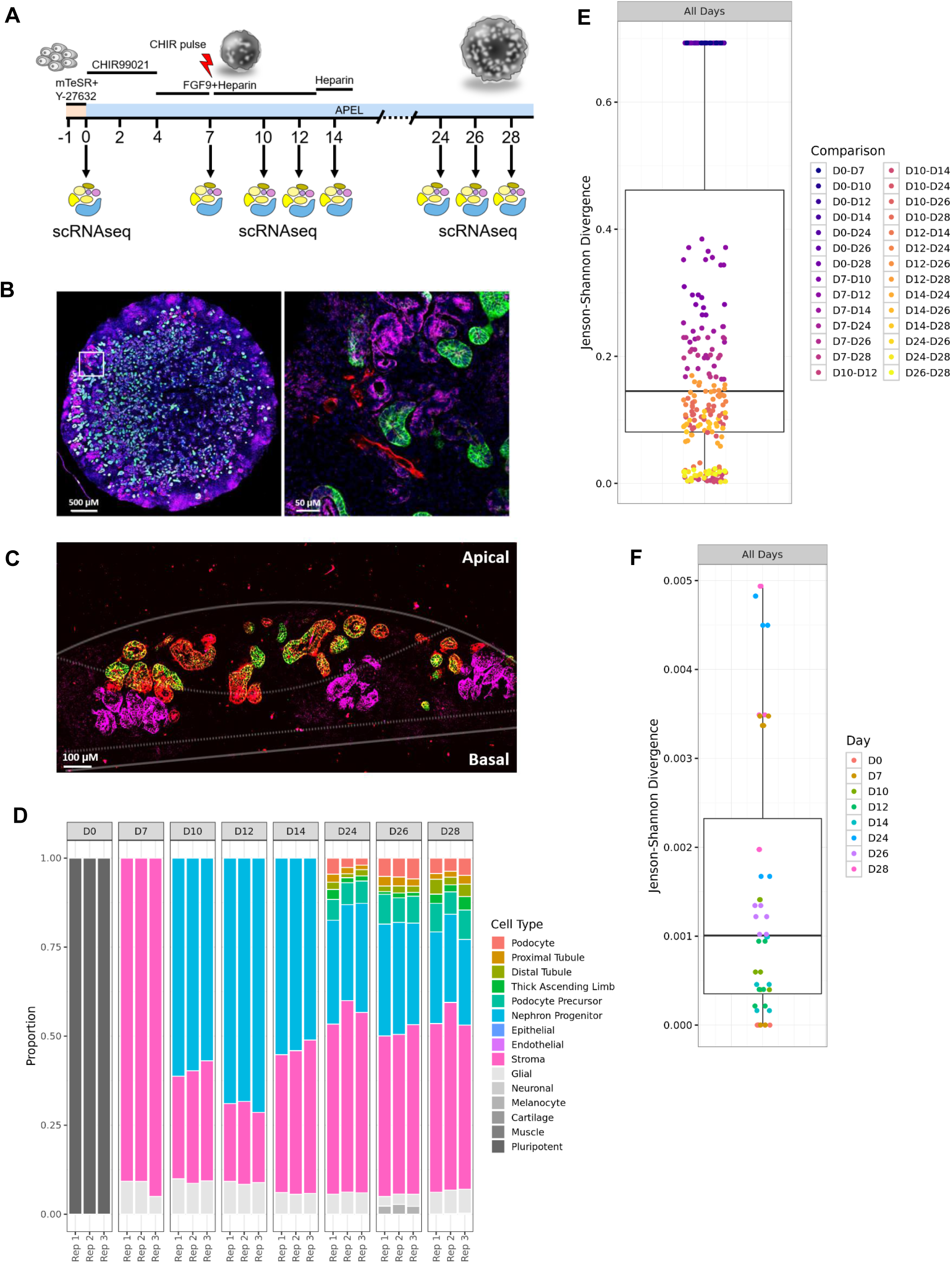
A high-throughput platform for human kidney organoid generation. (A) Schematic of organoid protocol with scRNA-Seq timepoints (B) Representative planar cross-section of a D28 *in vitro* organoid that expressing markers of podocytes, tubules, and endothelial cells (*left panel*). A zoomed in region of the cross-section showing endogenous human CD31/PECAM positive vessels developing *in vitro* amongst podocytes and tubules (*right panel*). Synaptopodin: magenta; CD31/PECAM1: red; E-cadherin: green; Hoechst: blue. (C) Representative cross-section of a D28 *in vitro* organoid demonstrating apical and basal nephron distribution of podocyte-containing glomeruli, proximal tubules, and distal tubules. Synaptopodin: magenta; LTL: red; E-cadherin: green. (D) Stacked bar graph of cell type proportions of *in vitro* organoid differentiation across time points and replicates reveal an increase in podocytes and tubular cells at later time points. (E) Comparison of cell type proportions across days using boxplots of Jenson-Shannon Divergence (JSD) indices show high similarity between time points. (F) Comparison of cell type proportions across replicates using boxplots of Jenson-Shannon Divergence (JSD) indices show high similarity between replicates of same time point.

### Transcriptional profiling reveals reproducibility of high-throughput organoids

The time period between day 7 and day 15 is critical for kidney organoid differentiation and variability (Subramanian et al., 2019). To determine whether the novel high-throughput protocol was compatible with appropriate differentiation and reproducibility, organoids were assessed by single cell profiling at day 0 (D0; iPSCs state), and salient intermediate time points including day 7 (D7), day 10 (D10), day 12 (D12), and day 14 (D14). We also profiled organoids on day 24 (D24), day 26 (D26), and day 28 (D28) *in vitro*. We successfully profiled 204,748 single cells from 24 organoids across all time points (Fig. 1A, D). iPSCs on D0 maintained a high level of pluripotency (Fig. 1D, Suppl. Fig. 1A) and matched published transcriptomic signatures (Subramanian et al., 2019). On D7, developing organoids expressed appropriate markers of mesodermal differentiation with actively proliferating cells (Kowalczyk et al., 2015) and little variability between replicates (Fig. 1D, Suppl. Fig. 1B). On D7, we recovered a small population of cells expressing epithelial cell makers, and by D10 nephron progenitor cells appeared, which presented the majority of cells through D14 (Fig. 1D, Suppl. Fig. 1C-E). At D24 through D28, we recovered podocytes and tubular epithelial cells (Fig. 1D, Suppl. Fig. 1F-H). Differences in cell type proportions were observed across timepoints (avg. JSD = 0.26, SD = 0.19, Fig. 1E, Suppl. Fig. 2A), while differences in proportions across replicates (Subramanian et al., 2019) were small (avg. JSD = 0.001, SD = 0.001, Fig. 1F, Suppl Fig. 2B). Thus, the majority of variations, which became first apparent at D10, were due to differences between stages of organoid differentiation and not due to variability between replicates in the same stage. We concluded that the differentiation of high-throughput kidney organoids, based on our scalable protocol, is highly reproducible.

**Figure 2.**
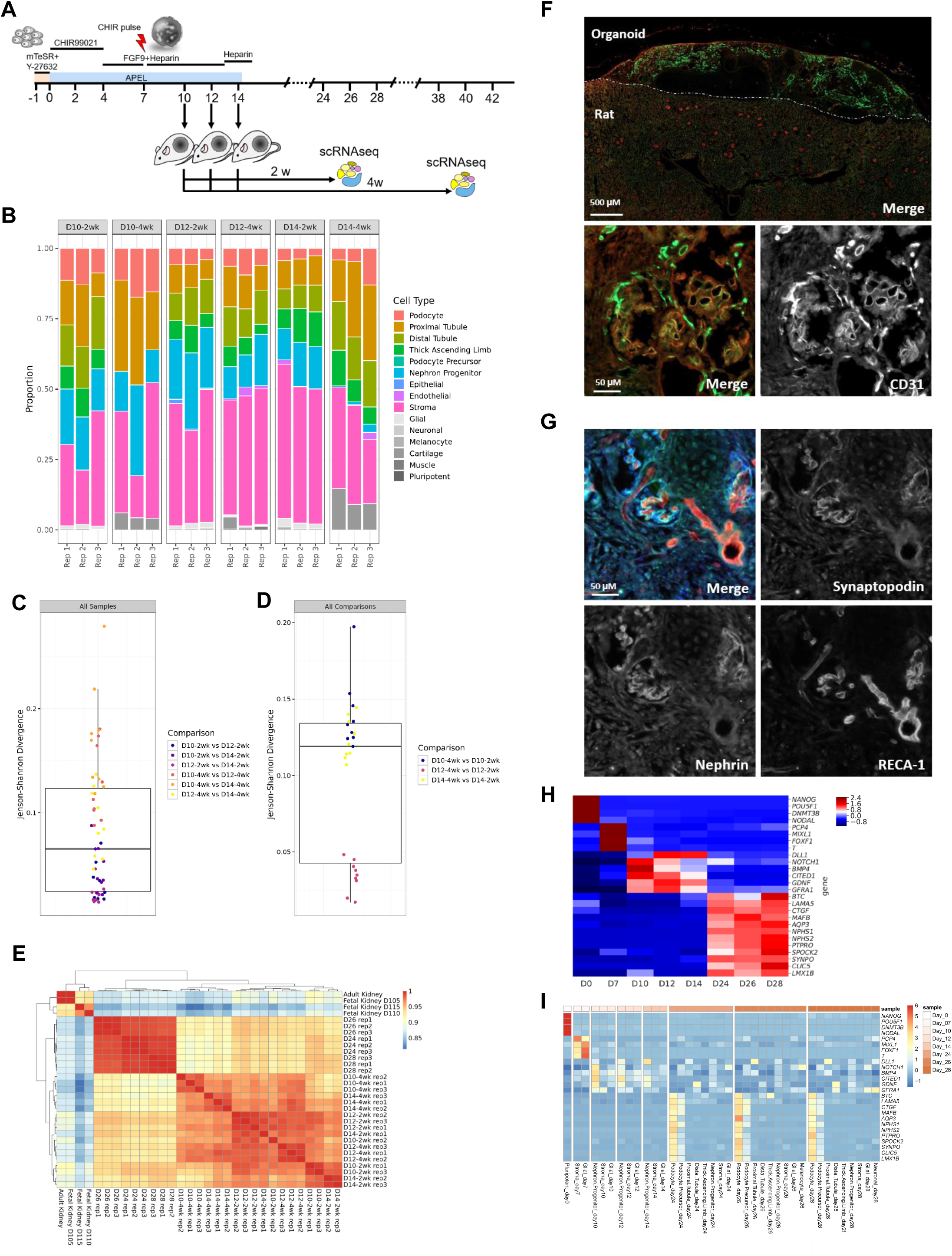
Organoid transplanted promotes differentiation and vascularization. (A) Schematic of organoid protocol with rat kidney capsule transplantation and scRNA-Seq timepoints (B) Stacked bar graph containing cell type proportions of *in vivo* maturation across time points and replicates show an increase in podocytes, tubular cells, endothelial cells, and off-target cell types with a longer period of *in vivo* maturation. (C) Comparison of cell type proportions across days using boxplot of Jenson-Shannon Divergence (JSD) indices reveals high similarity between different days. (D) Comparison of cell type proportions across weeks using boxplot of Jenson-Shannon Divergence (JSD) indices shows high similarity between different weeks. (E) Hierarchical clustering of Spearman correlations of gene expression profiles of podocytes from adult human samples, fetal human samples, *in vitro* organoids, and transplanted organoids. (F) Representative transplanted organoid showing CD31/PECAM expressing cells indicating single human endothelial cells and small vessel networks (Top panel) and a zoom representative showing CD31/PECAM cells in a organoid glomerulus (Bottom panels). RECA-1: red; CD31/PECAM1: green. (G)Representative transplanted organoid showing RECA-1 expressing rat derived vascularization of an organoid glomerulus. RECA-1: red; synaptopodin: green; nephrin: blue (H) NanoString gene expression (row z-score) over *in vitro* organoid maturation (D0 to D28) for selected genes. (I) Average gene expression scaled to row z-score within each scRNA-Seq cluster during *in vitro* organoid maturation (D0 to D28) for selected genes.

### Transplantation promotes organoid vascularization and maturation

Organoids *in vitro* are not perfused by the systemic circulation, thereby precluding PK/PD studies, an essential component of drug discovery. To address this issue, we adapted organoid transplantation protocols (Sharmin et al., 2016; Subramanian et al., 2019) from mice to rats, because rats are most commonly used for PK/PD studies (Fig. 2A). We grew organoids *in vitro* for 10, 12 or 14 days before inserting them under the kidney capsule of athymic rats, and further grew them for 2 or 4 weeks *in vivo* before harvesting the organoids for scRNA-Seq, NanoString, and imaging analysis (Fig. 2A). To determine the optimal time point for kidney organoid transplantation, we profiled 143,475 cells from 18 transplanted organoids by scRNA-Seq. Compared to *in vitro* growth, the proportion of differentiated podocytes and tubular epithelial cells increased with *in vivo* maturation (Fig. 2B, Suppl. Fig. 3A-F). A longer period of *in vivo* maturation resulted in a larger proportion of differentiated podocytes (0.098 vs 0.066, Χ2 = 461, p < 1e-15), proximal tubules (0.209 vs 0.112, Χ2 = 2436, p < 1e-15), distal tubules (excluding D10: 0.142 vs. 0.100, Χ2 = 158, p < 1e-15), and endothelial cells (0.008 vs 0.004, Χ2 = 89, p < 1e-15). When comparing 2 versus 4 weeks of *in vivo* growth, we noted more off-target cell types in the 4-week transplanted organoids (0.006 vs 0.002, Χ2 = 1520, p < 1e-15) (Fig. 2B, Suppl. Fig. 3A-F). Differences between timepoints were observed between transplant days (median JSD = 0.08, SD = 0.03, Fig. 2C, Suppl Fig. 4A) as well as between weeks of *in vivo* maturation (median JSD = 0.10, SD = 0.02, Fig. 2D, Suppl Fig. 4B). *In vivo* maturation for 2 weeks was more highly correlated with scRNA-Seq profiles from fetal and adult human kidney (Spearman’s ρ = 0.880, SD = 0.018) than *in vivo* maturation for 4 weeks (Spearman’s ρ = 0.858, SD = 0.019, *t* = 5.09, p = 2.8e-6, Fig. 2E). Additionally, within the distal tubule cluster, we observed a small population of cells expressing *AQP2* and *CALB1*, markers of collecting ducts (Subramanian et al., 2019) (Suppl. Fig. 5A-F). This subpopulation of collecting duct cells was most evident in D10 organoids after *in vivo* maturation for 2 weeks (Suppl. Fig. 5A-F). The transplanted organoids underwent vascularization by both rat RECA-1-expressing and human CD31/PECAM1-expressing endothelial cells (Fig. 2F-G, Suppl. Fig. 6A) The upregulation of human CD31/PECAM1 in transplanted organoids was verified by NanoString mRNA analysis and highest levels were found in D14 organoids grown *in vivo* for 4 weeks (Suppl. Fig. 6B). We also confirmed *AQP2* mRNA upregulation by NanoString (Suppl. Fig. 6C). Taken together, transplantation on D10 and *in vivo* maturation for 2 weeks was optimal for kidney organoid maturation, as reflected by the presence of all nephron segments, including collecting ducts, and the lowest number of off-target cells. Conversely, transplantation on D14 and *in vivo* maturation for 4 weeks was best for organoid vascularization, although this came at the price of acquiring more off-target cells.

### Development of a novel NanoString panel for rapid organoid QC

The single cell RNAseq (scRNA-Seq) time course provides a high-resolution analysis of cell proportions and their respective gene expressions during organoid differentiation. Even though scRNA-Seq is a powerful tool for organoid single cell profiling (Tanay and Regev, 2017), cost and time limit its use for routine organoid QC. NanoString is a multiplexed gene expression assay used to quantify absolute RNA counts (Geiss et al., 2008). Importantly, NanoString is a rapid, hands-off assay that can provide high quality data in 48 h. We used the scRNA-Seq profiles to develop a NanoString panel for rapid QC of iPSCs and organoid batches before selecting them for *in vitro* experiments or transplantation studies (Suppl. Fig. 7 and Suppl. Fig. 8). A set of 222 genes marking pluripotency, differentiation-regulating transcription factors, and cell-type specific genes (and 6 reference genes; Table 1) were used to interrogate mRNA expression levels during each timepoint previously analyzed by scRNA-Seq. To determine how closely NanoString and scRNA-Seq data matched, we extracted gene expression counts for all genes on the NanoString panel from the scRNA-Seq dataset, averaged by cell cluster. We compared these data to the NanoString RNA data and derived heatmaps of hierarchically clustered gene expression to group genes that vary similarly across developmental stages in both assays (Suppl. Fig. 7 and Suppl. Fig. 8). We selected a few representative genes to illustrate the similarity in expression patterns of NanoString (Fig. 2H, Suppl. 7) and scRNA-seq (Fig. 2I, Suppl. 8). A clear decrease of pluripotency genes was observed between D0 and D7 (Fig. 2H, I, Suppl. 7, 8). Expression of transcriptional programs important for kidney development such as Notch signaling (Sirin and Susztak, 2012) at D7 and D10-14 indicated that kidney differentiation had been properly initiated (Fig 2H, I, Suppl. 7, 8). The NanoString data revealed the presence of mature kidney cells (Fig. 2I, Suppl. 7). For example, we found, by scRNA-Seq, 12 genes that were highly expressed in mature podocytes but to a lesser degree in podocyte progenitors at D24 and D28 (Fig. 2I, Suppl. 8), and these results were confirmed by the NanoString analysis (Fig. 2H, Suppl. 7).The NanoString assay was performed on whole organoids and RNA expression counts thus represent a pool of all cell types present in the organoid. However, using cell type specific markers allows to obtain an estimate of the proportion of mature kidney cell types such as podocytes within kidney organoids. The same approach was used to derive NanoString marker panels for proximal, thick ascending limb, and distal tubular cells, as well as off-target cells such as glia, melanocytes and neurons (Suppl. Fig. 7 and Suppl. Fig. 8, Fig. 2I). Taken together, gene expression analysis by NanoString allowed for rapid, routine QC of iPSCs and organoids. This novel QC metric for organoids made it feasible to decide as early as D7 whether the quality was sufficient to continue organoid batch differentiation, or whether the quality was poor, necessitating that we discard the batch and start anew. Ultimately, using this novel approach, we noted significant benefits to both time and cost in developing a scalable, high-throughput organoid platform.

### Application of the organoid platform to clinically relevant scenarios

Having established QC metrics for the organoid platform, we turned to its functional validation for drug discovery. To this end, we conducted proof-of-concept studies with the calcineurin inhibitor CsA, which is clinically used to treat patients with proteinuric kidney diseases (Mathieson, 2008) in comparison with GFB-887, a novel potent, small molecule inhibitor of TRPC5 ion channels belonging to a family of subtype-selective TRPC5 inhibitors (Yu et al., 2019). Whole-cell patch clamp electrophysiology in human TRPC5-expressing HEK293 cells showed GFB-887-mediated inhibition of TRPC5 current (Fig. 3A). The previously described TRPC5 inhibitor tool compound ML204 served as positive control (Tian et al., 2010) (Fig. 3A). The IC50 of GFB-887 was 0.037 μM (Fig. 3B). By scRNA-seq we detected a few cells that expressed *TRPC5* mRNA. Before embarking on further functional studies, we confirmed the scRNA-Seq expression of *TRPC5* in iPSCs and kidney organoids by NanoString analysis. We detected a time-dependent upregulation of *TRPC5* mRNA expression during organoid differentiation and found highest *TRPC5* mRNA levels in D40 to D45 organoids (Fig. 3C). The presence of TRPC5 protein in organoids was confirmed by double labeling immunofluorescence microscopy with the podocyte marker synaptopodin (Mundel et al., 1997) (Fig. 3D, Suppl. Fig. 9).

**Figure 3.**
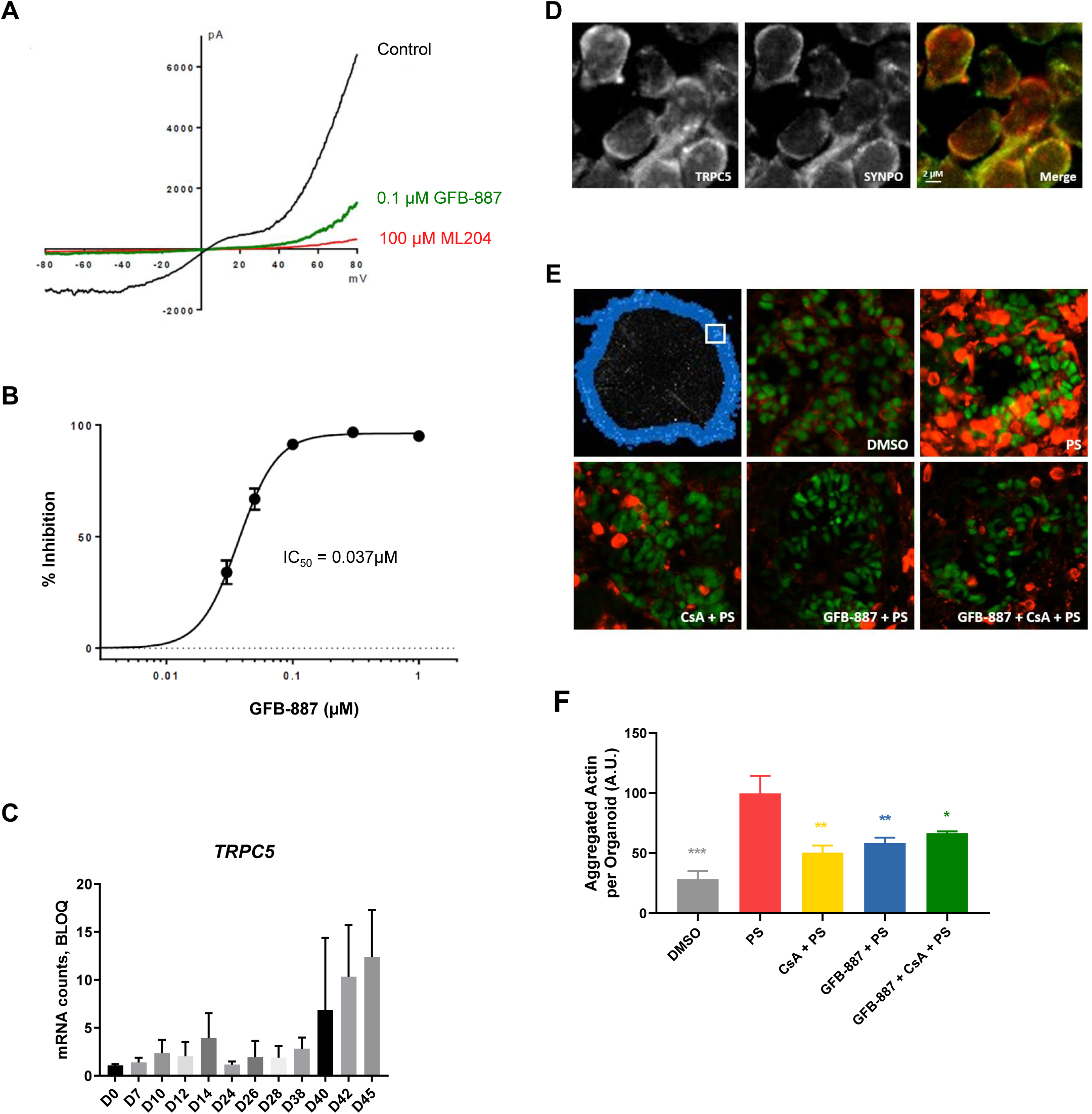
CsA and TRPC5 inhibition protects against podocyte injury in kidney organoids *in vitro*. (A) Representative current-voltage relationship recording during a 500 ms voltage ramp from -80 mV to +80 mV in the absence and presence of 0.1 μM GFB-887 or the reference TRPC5 inhibitor, 100 μM ML-204. (B) Concentration dependence of inhibition of human TRPC5 currents after a 3 min application of GFB-887 at +80 mV. Data points are mean ± SEM of 3-4 observations at each concentration. (C) Upregulation of *TRPC5* mRNA expression during organoid differentiation *in vitro* (D) Double labeling with synaptopodin reveals podocyte TRPC5 protein expression in human organoids (E) CsA and GFB-887 protects against PS induced podocyte injury in *in vitro* D28 organoids. Representative PS injury mask for an organoid is depicted in blue and the inset box indicates and area in the region that can be used for identification of injured podocytes for quantitation (*top left panel*). Representative glomeruli identified for podocyte injury quantitation of aggregated actin for treatment with vehicle DMSO, PS alone, CsA + PS, GFB-887 + PS, and the combination of CsA and GFB-887 + PS. Synaptopodin: green; Phalloidin: red. (F) Quantification of PS induced actin aggregation. GFB-887 and CsA are non-additive, consistent with shared mechanism of action. DMSO vs. PS, p<0.0001; CsA + PS vs. PS, p = 0.0021; GFB-887 + PS vs. PS, p = 0.0087, CsA + GFB-887 + PS vs. PS, p = 0.0397. Data show mean ± SEM.

**Figure 4.**
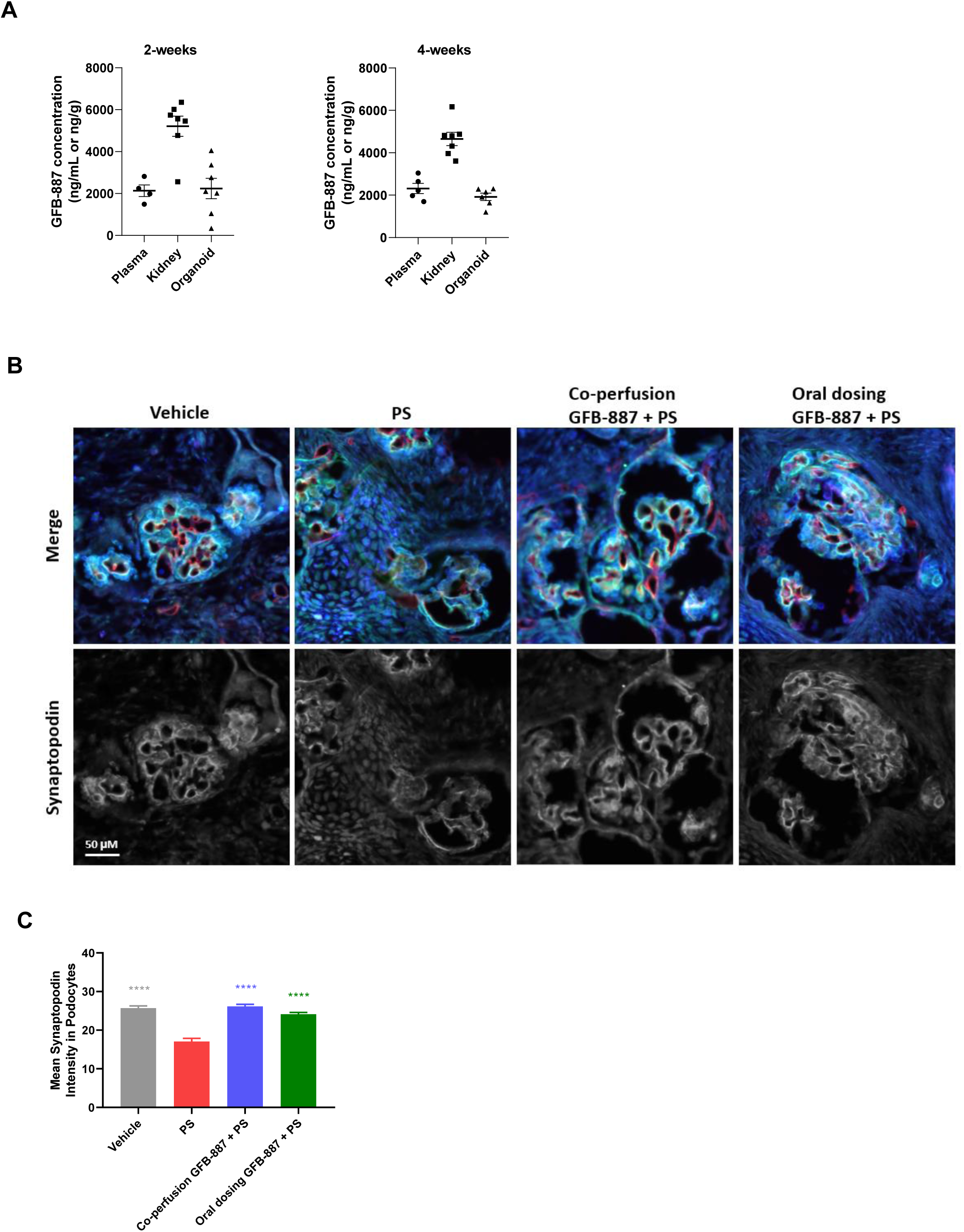
TRPC5 inhibition protects against podocyte injury in human kidney organoids *in vivo*. (A) Oral dosing of GFB-887 in rats results in drug exposure of transplanted, perfused human kidney organoids equivalent to rat plasma levels. Athymic nude, male rats were implanted with day 14 early differentiating organoids under the sub-capsular space. Transplanted rats were grown for 2 (*left panel*) or 4 (*right panel*) weeks and orally dosed with 10 mg/kg GFB-887 for 3 consecutive days prior to plasma, kidney and organoid collection. Data show mean ± SEM of GFB-887 exposure from at least 4 independent measurements suggest that transplanted human kidney organoids achieve maximal vascularization as well as functional connectivity to host vasculature in 2 weeks. (B) Super resolution imaging reveals PS-induced loss of synaptopodin in transplanted human organoids, which can be prevent by orally dosing with GFB-887: HBSS (Hank’s Balanced Salt solution serving as vehicle control), PS, PS + GFB-887. (C) Quantification of PS induced podocyte injury and protection by CsA or GBF-887. Synaptopodin mean intensity in podocytes was quantified from human organoids from transplant experiments for vehicle, PS, co-perfusion GFB-887 + PS, and oral dosing of GFB-887 + PS (Data show mean ± SEM; for all treatment conditions vs. PS p<0.0001).

Focal segmental glomerulosclerosis (FSGS) is a disorder of podocytes with high likelihood of progression to kidney failure (D’Agati et al., 2011). In animal models of FSGS, inhibition of TRPC5 channel activity with tool compounds protects against proteinuria and podocyte loss (Yu et al., 2019; Zhou et al., 2017). Based on preclinical data in cell culture and animal models with tool compounds (Schaldecker et al., 2013), we assessed the protective effects of CsA and GFB-887 on protamine sulfate (PS)-induced podocyte cytoskeletal injury (Buvall et al., 2017; Schaldecker et al., 2013). Similar to its effect on mouse podocytes grown in a monolayer *in vitro* (Buvall et al., 2017), PS induced aggregation of actin fibers and increased phalloidin labeling in organoid podocytes, and both CsA and GFB-887 protected from PS-driven actin aggregation (Fig. 3E). For quantitative analysis, we measured the changes in phalloidin labeling in sections of entire organoids (blue ring in Fig. 3E). The quantitative analysis confirmed the protective effect of CsA and GFB-887 (Fig. 3F). In keeping with the observation that TRPC5 is the ion channel required for calcineurin activation in podocytes (Tian et al., 2010), and therefore the two molecules are targeting the same signaling pathway, the efficacy of CsA and GFB-887 were comparable and no additive effect was noted (Fig. 3E, F).

### GFB-887 protects against podocyte injury in transplanted human organoids

Finally, we conducted PK/PD studies in transplanted organoids in rats to evaluate the effect of GFB-887 on PS-induced podocyte cytoskeletal injury *in vivo* in human transplanted organoids. We found that oral dosing of GFB-887 (10 mg/kg) to rats carrying D14 organoids transplanted for 2 weeks or 4 weeks resulted in measurable drug exposure, not only in rat plasma and rat kidneys, but also in the transplanted human organoids (Fig. 4A). Next we assessed PS-induced podocyte injury in transplanted organoids. Historically, the readout of PS studies had been transmission electron microscopy of kidney tissue (Kerjaschki D, 1978; Schaldecker et al., 2013; Seiler et al., 1975), which is labor-intensive and time consuming. To address this, we developed a novel, efficient, approach for the quantification of PS-induced loss of synaptopodin protein abundance (Buvall et al., 2017; Schaldecker et al., 2013) that combines fluorescence and super-resolution microscopy (Suleiman et al., 2017). We found that PS caused a significant reduction in synaptopodin abundance in organoids (Fig. 4B). Co-perfusion of 3 μM GFB-887 with PS or oral dosing of GFB-887 (10 mg/kg) prior to PS perfusion, preserved the abundance of synaptopodin at levels similar to those found in saline-perfused control organoids (Fig. 4B), and the quantitative analysis confirmed these results (Fig. 4C). Collectively, these data show that inhibition of TRPC5 by GFB-887 effectively protects human podocytes in kidney organoids, both *in vitro* and *in vivo*. Thus, we have successfully established a translational human kidney organoid platform which can be used to evaluate the pharmacodynamic efficacy of known and novel compounds with therapeutic potential in a human system prior to clinical development.

## Discussion

In this study, we addressed prior concerns about the applicability and scalability of human organoids by building a high-throughput, quality controlled organoid platform for preclinical studies in a human system. As a proof of concept, we used the platform to illuminate the human relevance of the TRPC5 inhibitor GFB-887, a novel drug candidate now being tested in the clinic (NCT03970122). This study has led to several important observations.

First, we developed a high-throughput (> 500 organoids/week) platform for differentiated, three-dimensional kidney organoids with reproducible quality suitable for drug discovery. Monitoring platform reproducibility at single cell resolution ensured standardization of the data. Based on insights from scRNA-Seq, we developed a NanoString panel for rapid and inexpensive QC of iPSC maintenance and organoid differentiation *in vitro* before and after transplantation. This is of particular importance given the complexity, timelines and costs of PK/PD studies with transplanted organoids.

Second, an important outcome of our studies was the observation that transplanted organoids developed a vasculature that was functionally connected to the rat host circulation, capable of successfully delivering compounds to this human system, an essential prerequisite for PK/PD studies. The combination of *in vitro* mechanistic studies in organoids with *in vivo* PK/PD studies in perfused organoids provides preclinical confidence in the human relevance of specific targets and compounds previously vetted only *in vitro* and/or in animal models. As a proof of concept, we established the human relevance of two compounds: a drug already used in the clinic (CsA) and a novel TRPC5 channel blocker (GFB-887), in advance of testing it in patients.

Third, the current study has highlighted the relevance of kidney organoids for glomerular diseases characterized by podocyte injury (Greka and Mundel, 2012). Future studies may build upon this work to employ patient-iPSC-derived organoids and to model more complex diseases such as diabetic kidney disease, which affects not only podocytes but also other kidney cells. Another open question for further investigation relates to the role of ureteric bud containing organoids. Our studies have revealed that *in vivo* maturation of transplanted D10 organoids can lead to the appearance of collecting duct cells. A recent study showed how co-culturing iPSC-derived ureteric bud and nephron progenitor cells induces collecting duct-containing, higher-order kidney organoids *in vitro* (Taguchi and Nishinakamura, 2017). Future studies are needed to determine whether these higher-order organoids confer advantages that merit their implementation in preclinical drug evaluation, in particular for the assessment of drugs that target the collecting duct in diseases such as polycystic kidney disease (Grantham, 2008).

In conclusion, the integrated human organoid platform described here markedly enriches preclinical drug evaluation by lending direct human relevance to animal studies before embarking on clinical trials. More generally, our data support the idea that, if properly maintained and quality controlled, human iPSC-derived organoid systems are likely to become an integral component of drug discovery programs in several tissues and organs, including, for example, the kidney, gut, retina, and brain. We are therefore at the dawn of a new era in drug discovery, where confidence in bringing drugs to the clinic may be derived, at least in part, from human organoid platforms.

## Acknowledgements

We thank Evan Murray for technical support, the Whitehead Institute Genome Technology Core for scRNA-Seq library preparation and sequencing, the Harvard Center for Biological Imaging (HCBI) for help with imaging, Brendan Fish and the Biomere team for conducting the organoid transplantation studies, and Icagen for conducting the electrophysiology studies.

## Materials and Methods

### Maintenance of human iPSCs

Human episomal iPSCs (ThermoFisher, Waltham, MA) were maintained on hESC-qualified matrigel (Corning, Corning NY) coated T25 flasks in mTeSR1 medium (StemCell Technologies, Vancouver, Canada) and passaged at 80-85 % confluency using Gentle Cell Dissociation Reagent (StemCell Technologies). For evaluation of pluripotency by flow cytometry, iPSCs were dissociated with Accutase (StemCell Technologies), washed with DPBS and fixed in 4 % PFA for 20 min at RT and rinsed with DPBS. The cells were then incubated for 1 h in blocking buffer (DPBS containing 0.3 % of Triton X-100, 1 % BSA, and 5 % donkey serum) and for 1 h at 4 0C with a combination of conjugated antibodies to surface proteins TRA-1-60 (StemCell Technologies, 60064PE), TRA-1-81(StemCell Technologies, 60065AZ), key transcription factors OCT3/4 (SantaCruz Biotechnology, sc-5279), NANOG (ThermoFisher Scientific, 53-5761-80) and a negative surface marker SSEA1(StemCell Technologies, 60060PE). Cells were washed with DPBS followed by data acquisition on a SONY SH800 flow cytometer.

### High-throughput kidney organoid generation

A high-throughput (> 500 organoids/week) protocol for human kidney organoid generation was developed based on a published protocol (Takasato et al., 2016a; Takasato et al., 2016b) with modifications described below (Fig. 1A). iPSCs reaching 80 to 85 % confluency were dispersed into single cell suspensions using Accutase and 275,000 cells per flask were plated in hESC-qualified, matrigel-coated flasks with mTeSR1 media plus Y-27632 (Tocris 1254, Bristol, United Kingdom). On the next day, the medium was changed to STEMdiff APEL2 medium (StemCell Technologies) containing 10 μM CHIR99021 (Tocris 4423). After 48 hours, the medium was changed to STEMdif APEL2 supplemented with 8 μM CHIR99021. Subsequently, the medium was changed to STEMdif APEL 2 with 200 ng/mL FGF9 (R&D Systems, Minneapolis, MN) and 1 mg/mL heparin (StemCell Technologies) and replaced every other day. On day 7, the cells were passaged using Accutase and pelleted by centrifugation at 500,00 cells per pellet. The pellets were placed onto transwell plates and STEMdiff APEL2 with 5 μM CHIR99021 was added for 1 hour before changing to STEMdiffAPEL2 with 200 ng/mL FGF9 and 1 mg/mL heparin. The medium was changed every other day using STEMdiff APEL2 with 20 ng/mL FGF9 and 1 mg/mL heparin for the next 4 days. On day 13, the medium was changed to STEMdiff APEL2 with 1 mg/mL heparin. Thereafter, the organoids were grown in STEMdiff APEL and medium was changed every other day.

### Organoid transplantation

Organoids were transplanted to 3-4 weeks old athymic nude rats as per approved IACUC protocol. The rats were anesthetized, kidneys were exposed after incisions in the abdominal wall, and after making an incision in the kidney capsule, a small space was created under the kidney capsule to host the organoids. Kidney organoids grown for 10, 12 or 14 days *in vitro* were aspirated into the tip of an 18-24G catheter (Instech, Plymouth, PA) and placed into the subcapsular space. After organoid implantation, the abdominal wall and skin were closed, and the animals were allowed to recover. At 2 or 4 weeks after transplantation, the organoids were harvested for scRNA-Seq and NanoString mRNA analysis. In addition, a set of organoids was collected for the analysis of graft tissue vascularization and functional connectivity in the PK/PD studies. Organoid differentiation was also evaluated by flow cytometry of two podocyte surface proteins, nephrin (Kestila et al., 1998) and podocalyxin (Kerjaschki et al., 1984). Cells from dissociated organoids were washed with DPBS, blocked in 1 % mouse serum for 30 min at 4 0C, incubated with anti-nephrin primary antibody (R&D Systems, AF4269) and APC-conjugated anti-podocalyxin antibodies (R&D Systems, FAB1658A) diluted in sorting buffer (HBSS containing 1% BSA and 0.035 % NaHCO_3_). The cells were further incubated for 10 min in Alexa Fluor 488-conjugated donkey anti-sheep LgG (H+L) antibody (ThermoFisher Scientific, A-11015), washed and resuspended in sorting buffer. Unstained cells were utilized to establish background fluorescence. Data acquisition and analysis were done on a SONY SH800 flow cytometer. Mean fluorescence in live cells was determined for each sample and is presented after subtraction of background fluorescence.

### Gene expression analysis by scRNA-Seq

Organoids were rinsed with DPBS, collected into 1.5 ml conical tube containing 500 μl TryplE Select and incubated for 5 min at 37 0C, passed 3 to 4 times through a 27G needle, and incubated for 5 min at 37 0C after adding additional 500 µl of TryplE Select. The organoids were dissociated into single cell suspensions by passing 3 to 4 times through a 30G needle, followed by centrifugation at 600 x g for 5 min. Supernatants were discarded, and cell pellets were resuspended in DPBS and filtered through a 40 μM cell strainer. Cell numbers were determined on a Cellometer cell counter and adjusted to 1000 cells/μL in DPBS. The cells were placed on ice, transferred to the RNA-Seq facility, introduced to droplet-based (10X) lysis and amplification, and sequenced on Illumina HiSeq. For cells from *in vitro* organoids, sequences were aligned to human reference (hg19) and, for cells from transplanted organoids, combined human and rat reference (Rnor 6.0) genomes. Replicates within time points were grouped, and a preliminary unbiased clustering was performed on the top 50 principal components (PCs) of gene expression. Cells with high ribosomal gene content and low read count were considered poor-quality cells and excluded from further analyses. The remaining cells were filtered by the number of recovered genes and mitochondrial gene content. In total, high-quality libraries were generated for 348,223 cells that had passed QC. To infer cell type populations within each time point, we used unbiased clustering of the top 50 PCs of residual gene expression after regressing out replicate, total gene count, mitochondrial gene content, and cell cycle score. We assigned cell type labels based on upregulation of marker genes in the focal cluster as determined by a Wilcoxon Rank Sum test in Seurat 3.1 in R 3.6.1. Divergences among timepoints and replicates were compared with Jansen-Shannon Divergence (JSD) indices (Subramanian et al., 2019). Cell type proportions were tested across conditions using chi square tests of independence. To compare podocyte maturation across timepoints and with data from fetal and adult human kidney samples (Menon et al., 2018; Wu et al., 2018), we calculated Spearman correlations between gene expression profiles and performed hierarchical clustering based on Euclidean distance between samples in R 3.6.1. (Phipson et al., 2019)

### RNA extraction and quantification

Total RNA was extracted from organoids or dissociated organoid single cell populations using the PureLink RNA Mini Kit (Life Technologies, Carlsbad, CA) following the manufacturer’s instructions. 200 μl lysis buffer were used per organoid for homogenization at 12,000 x g for 2 min in a QIAshredder Column (Qiagen, Hilden, Germany) after lysis and prior to addition of 70% ethanol. Total RNA was eluted with 50 μl RNase-free water and stored at -80 °C. Total RNA was quantified in a NanoDrop™ 2000c Spectrophotometer (Thermo Scientific).

### NanoString gene expression analysis

100 ng of total RNA in 5 μl of RNase-free water were subjected to NanoString nCounter Elements workflow, according to the nCounter Elements XT Assay User Manual (NanoString Technologies, Seattle, WA). mRNA expression of 228 genes was assessed using nSolver 4.0 Software (NanoString Technologies). The gene panel (Table 1) included pluripotency markers (e.g. *NANOG, POU5F1, SOX2*), kidney-specific markers (early development: *SIX2, LHX1, OSR1*; mature podocytes: *SYNPO, PODXL, NPHS1, NPHS2*), as well as markers of off-target cell types (e.g. *CRABP1, HES6, PMEL*). Gene expression levels were normalized using six reference genes (*ABCF1, GUSB, HPRT1, LDHA, POLR1B, and RPLP0*), which were included in the gene panel. Heatmaps were generated using the Python Seaborn package and displayed as hierarchical clustering (rows) of z-score transformed expression values. For comparison of NanoString data to scRNAseq data normalized, log-transformed counts for the 228 genes on the NanoString panel were extracted and averaged across each cluster (Seurat function “AverageExpression”). 18 genes that were not detected at any developmental stage were dropped. Heatmaps were generated using R package ‘pheatmap’. Row z-scaled gene expression matrix was clustered (hierarchically) using Euclidian distance metric.

### Electrophysiology

Manual patch-clamp electrophysiology was performed in the whole-cell configuration in HEK293 cells stably overexpressing human TRPC5 according to published protocols, using the identical external and internal solutions as previously reported (Tian et al., 2010). Once in the whole cell configuration, cells were held at 0 mV and TRPC5 current was measured by an initial step to -80 mV held for 50 ms, followed by a ramp protocol from -80 mV to +80 mV at a rate of 1 mV/ms, then a step at +80 mV for 50 ms before returning to the holding potential. The voltage protocol was repeated at a frequency of 1 Hz. Mean inward and outward currents were determined at -80 mV and +80 mV, respectively. GFB-887 and ML204 were delivered to cells diluted at final concentration in external buffer through a gravity perfusion system (Tian et al., 2010).

### Compound treatment of organoids *in vitro*

D24 to D28 *in vitro* organoids were used for CsA and GFB-887 compound treatment. CsA was diluted from a 20 mM stock in DMSO to final concentrations of 20 μM, 2 μM, 0.2 μM, or 0.02 μM in STEMdiff APEL2, GFB-887 from a 10 mM stock in DMSO to final concentrations of 10 μM, 1 μM, 0.1 μM or 0.01 μM in STEMdiff APEL2. Compounds stability in solution was analyzed using Waters Acquity UPLC. 1 ml of GFB-887 containing medium was added to the apical side of the transwell insert and 2 ml of GFB-887 containing medium were added to the basolateral side of the transwell insert. Organoids were pre-treated with compounds separately or in combination for 1 h before exposure to protamine sulfate at 300 μg/ml (Sigma) for an additional 1, 6, or 24 h.

### Immunofluorescence microscopy

Organoids grown *in vitro* or transplanted organoids with adjacent rat kidneys were rinsed with DPBS and fixed with 4% PFA at RT for 30 min, washed twice with DPBS for 5 min at RT, and incubated in 30 % sucrose at 4 °C for 2 to 3 days. After freezing the samples in cryoblocks using OCT (VWR, Radnor, PA), cryosectioning was done on a Leica CM1950 cryostat. Tissue sections were collected at 10 µm thickness, placed on glass slides, treated with 0.1% Triton-X 100, and blocked in 10% donkey or goat serum. After incubated with primary antibodies overnight at 4 °C, sections were washed with DPBS and incubated with secondary antibodies for 1 h at RT. Primary antibodies included: CD31/PECAM-1 (Bethyl Laboratories, IHC-00055), RECA-1 (BioRad, MCA970GA), synaptopodin (Progen, GP94-N), E-cadherin (R&D, AF648), TRPC5 (UC Davic/NIH NeuroMAB Facility, 75-104), TRPC5 (Alomon Labs, ACC-020). Secondary antibodies include: Alexa Fluor® 647 Donkey anti-Rat, Alexa Fluor® 555 Donkey anti-Mouse, Alexa Fluor® 555 Donkey anti-Rabbit, Alexa Fluor® 488 Donkey anti-Rabbit, Alexa Fluor® 488 Donkey anti-Goat (ThermoFisher Scientific), Alexa Fluor® 647 Donkey anti-Guinea Pig (Jackson Lab), Alexa Fluor® 405 Donkey anti-Goat (Abcam), Alexa Fluor® 750 Donkey anti-Goat (Abcam) were used at 1:300 for 1 h at RT, Hoechst was used at 1:10000 and phalloidin at 1:1000 for 30 min at RT. Fluorescence microscopy images were captured on a digital Axio Scan.Z1 Slide Scanner (Carl Zeiss White Plains, NY) or a superresolution confocal LSM 880 AiryScan microscope (Carl Zeiss). Zen5 software powered by Arivis was used for image acquisition and processing.

### Quantification of podocyte injury in organoids *in vitro*

For semi-automated image analysis with Amira for Cell Biology imaging software, the DAPI channel was used to identify cells and to create an organoid mask, the phalloidin channel to identify penetration depth of PS injury and to create a mask for region of injury quantitation, and the synaptopodin channel to identify injured podocytes. Using overlays of the DAPI channel mask to limit detection to organoid tissue on a slide, the phalloidin channel mask to limit detection to the region of injury, and the synaptopodin channel masks to limit detection to the regions containing podocytes, we used Amira to provide quantitative readouts of average phalloidin intensity over podocyte surface areas. GraphPad Prism 8.0.1 was used to graph the quantitative readout of aggregated actin and perform ordinary one-way ANOVA for multiple comparisons.

### PK studies in rats carrying transplanted organoids

For the PK studies, 5 to 6-week old athymic nude rats carrying D14 transplanted organoids were used. 10 mg/kg GFB-887 formulated in vehicle solution (Solutol HS-15 (Sigma)/vitamin E-TPGS (PMC Isochem, France)/PEG300 (Sigma)) or vehicle alone were dosed orally once a day for three consecutive days. For direct administration of GFB-887 or vehicle into the renal artery, a thoracotomy, according to approved IACUC protocol, was performed followed by clamping the supra- and infrarenal abdominal aorta below the diaphragm. After perfusion with HBSS at 9 ml/min for 5 min through a fine catheter (Exel, Santa Ana, CA) inserted into the clamped aorta, 3 μM GFB-887 in HBSS with 0.3% DMSO or HBSS with 0.3% DMSO were perfused for 15 minutes into the rat kidney/organoid pair. A small incision in the inferior vena-cava allowed drainage of the perfusion solutions. At the end of the perfusion, organoids and host kidneys were excised together, rinsed with ice-cold DPBS (Sigma), snap frozen in liquid nitrogen, and stored at -80 °C until further use. Blood samples (100-200 μl) were drawn 4 h post oral dosing on days 1 and 2 and at 0.5, 1, 2 and 4 h post dosing on day 3 and collected on ice in microcentrifuge tubes containing K2-EDTA (Fisher Scientific). Within 30 min after collection, blood samples were processed to plasma by centrifugation at 10,000 RPM at 4 °C. Plasma samples were snap frozen in liquid nitrogen and stored at -80 °C. GFB-887 concentrations in organoids, rat kidneys and plasma samples were measured by LC-MS at RMI Laboratories (North Wales, PA). Organoid and kidney sample processing involved initial homogenization with 0.5 mm Zirconium beads for 45 sec at 4,000 cycles/min in 500 to 1000 μl 50:50 ACN/water mixture containing 0.1 μg/ml internal standard and 0.1% FA. Supernatants were spun at 13,000 RPM for 5 min and 100 μl supernatant were mixed with 100 μl of 0.1% FA solution prior to LC/MS-MS analysis. Full scan LC-MS/MS data were analyzed with Quanlynx software for internal standards and GFB-887. The data were quantified using a matrix specific 6-point calibration curve spiked with GFB-887.

### PD studies in rats carrying transplanted organoids

PS-induced podocyte foot process effacement (Kerjaschki D, 1978) was assessed in rats carrying D14 organoids for 4 weeks. After clamping of the renal artery and excision of one organoid/kidney pair for PK studies, vascular perfusion of the contralateral kidney/organoid pair at a flow rate of 9ml/min was done as follows: i) vehicle group: 5 min HBSS, 30 min 0.3% DMSO, 5 min HBSS, 5 min fixative (4% PFA in DPBS); ii) PS group: 5 min HBSS, 15 min 0.3% DMSO, 15 minutes 2mg/ml PS in 0.3% DMSO; 5 min HBSS, 5 min fixative; iii) PS + oral GFB-887 dosing group: 5 min HBSS/15 min 2mg/ml PS in 0.3% DMSO/5 minutes HBSS washout/5 minutes fixative; iv) renal artery GFB-887 dosing group: 5 min HBSS, 15 min 2mg/ml PS and 3 μM GFB-887 in 0.3% DMSO, 5 min HBSS, 5 min fixative. After fixation, organoids and host kidneys were excised together, rinsed with ice-cold DPBS, trimmed and immersion fixed in 4 % PFA for 24 to 48 h at 4 °C. Samples were transferred to 30% sucrose, and stored at 4 °C.

### Quantification of podocyte injury *in* transplanted organoids and adjacent rat tissue

D14 organoids transplanted for 4 weeks and the adjacent endogenous rat kidneys were processed for immunofluorescence using Fiji/ImageJ 1.52g (Open source NIH imaging software). Staining for nephrin and synaptopodin was used to identify podocytes in organoid glomeruli. Using overlays of the nephrin channel mask to limit detection to podocytes, Fiji/Image J was used to quantitative readouts of synaptopodin average intensity over surface area within nephrin-positive masked areas. GraphPad Prism was used to graph the quantitative readout of synaptopodin intensity and perform ordinary one-way ANOVA for multiple comparisons. One-way ANOVA followed by Dunnett’s multiple comparisons test was performed using GraphPad Prism.

## Supplementary Figures and Tables

**Supplementary Figure 1.**
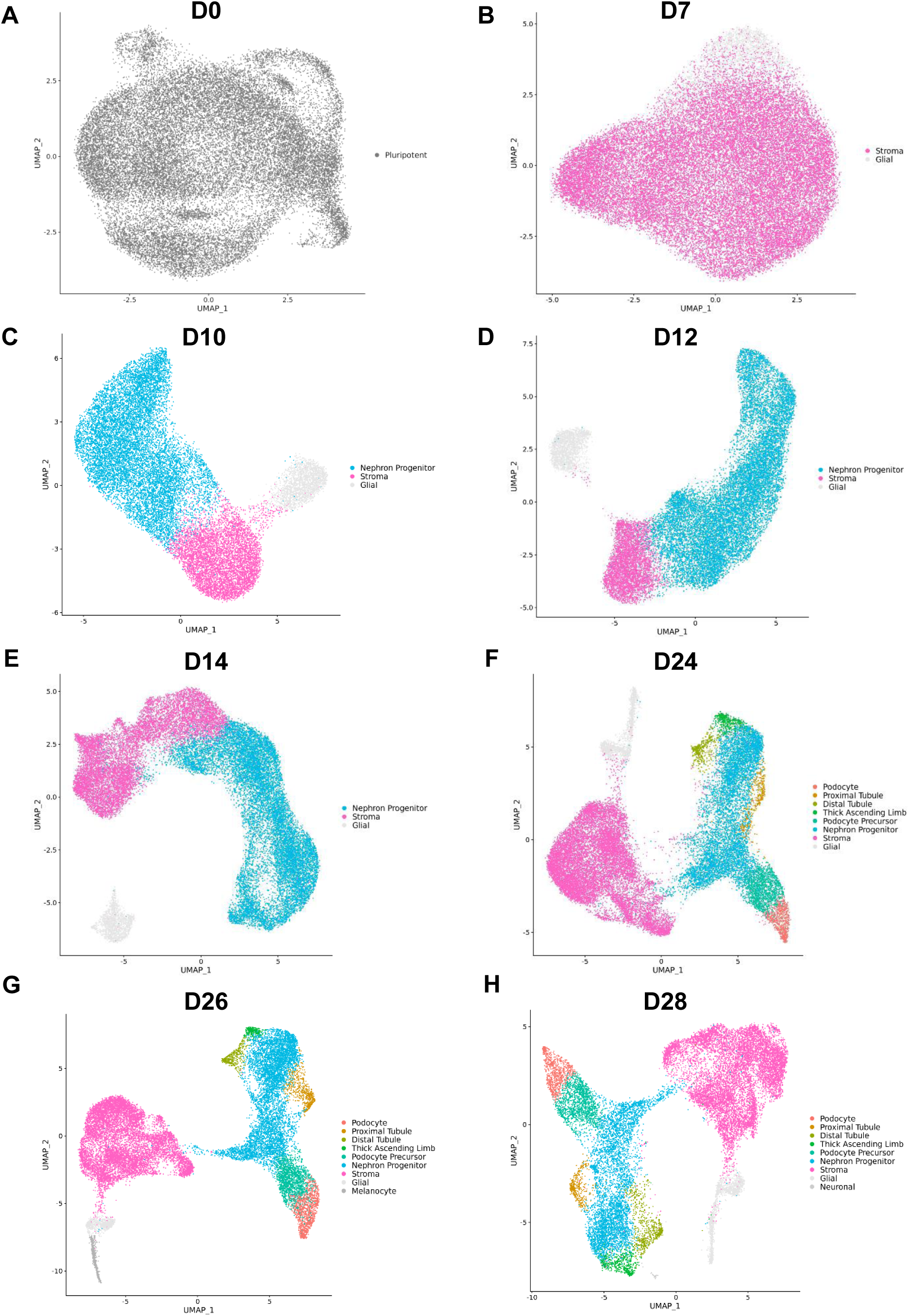
Two-dimensional representation of gene expression (Uniform Manifold Approximation and Projection, UMAP) of cell populations for *in vitro* timepoints across replicates. (A) On day 0 (D0) all cells are pluripotent. (B) On day 7 (D7) clusters of stroma and glial cells are present. (C-E) On day 10 (D10), day 12 (D12), and day 14 (D14), the majority of cells are nephron progenitors. (F) On day 24 (D24), clusters of podocytes and tubular cells are observed. (G) On day 26 (D26), in addition to clusters of podocytes, tubular cells, off-target cell types (melanocytes) are found. (H) On day 28 (D28) increased numbers of podocytes, tubular cells, and off-target cell types (neuronal) are present.

**Supplementary Figure 2.**
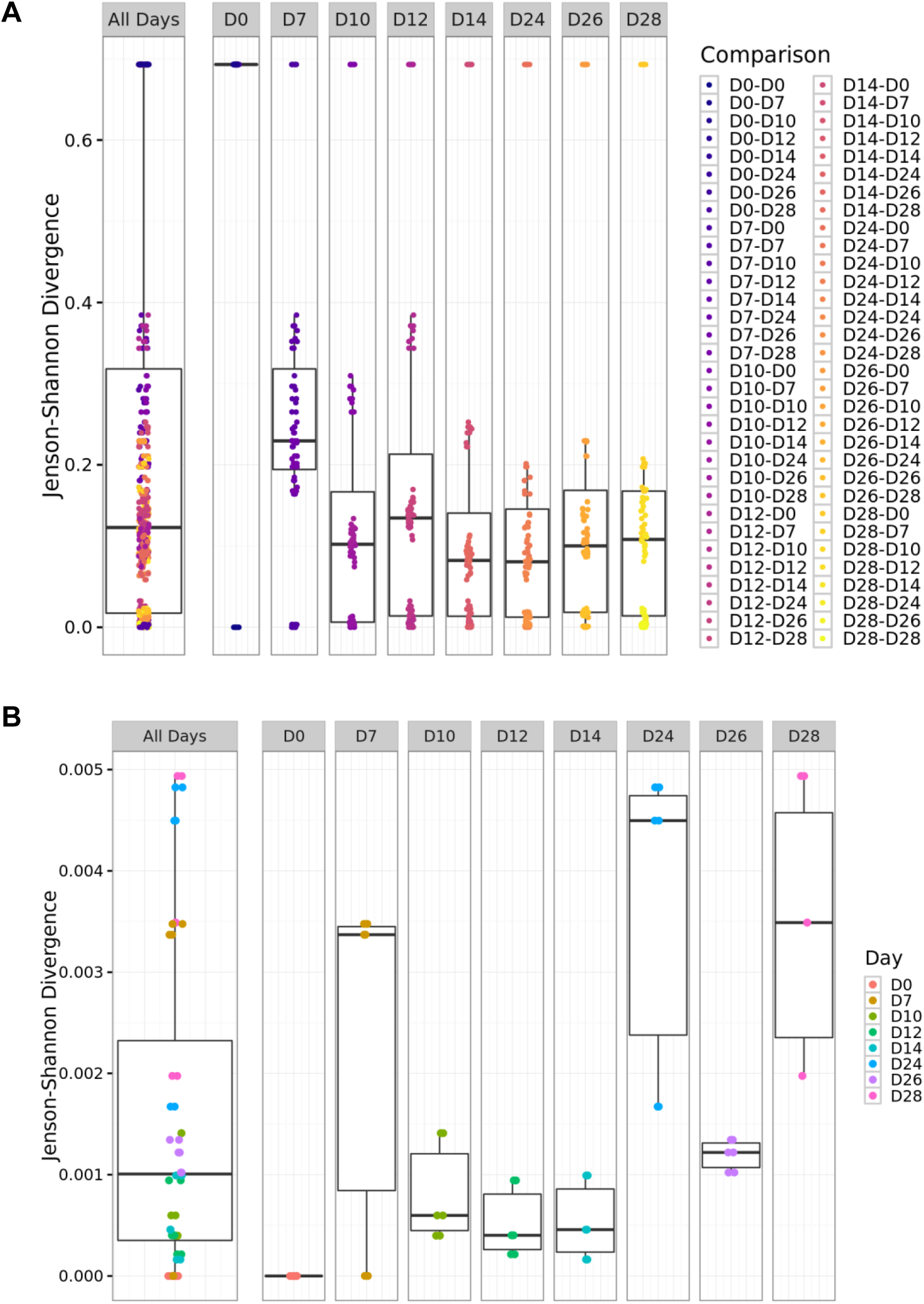
Comparison of cell type proportions for all *in vitro* timepoints using boxplots of Jenson-Shannon Divergence (JSD) indices. (A) Comparisons across time points shows increasing similarity with time. (B) Comparisons across replicates reveals increasing similarity with time.

**Supplementary Figure 3.**
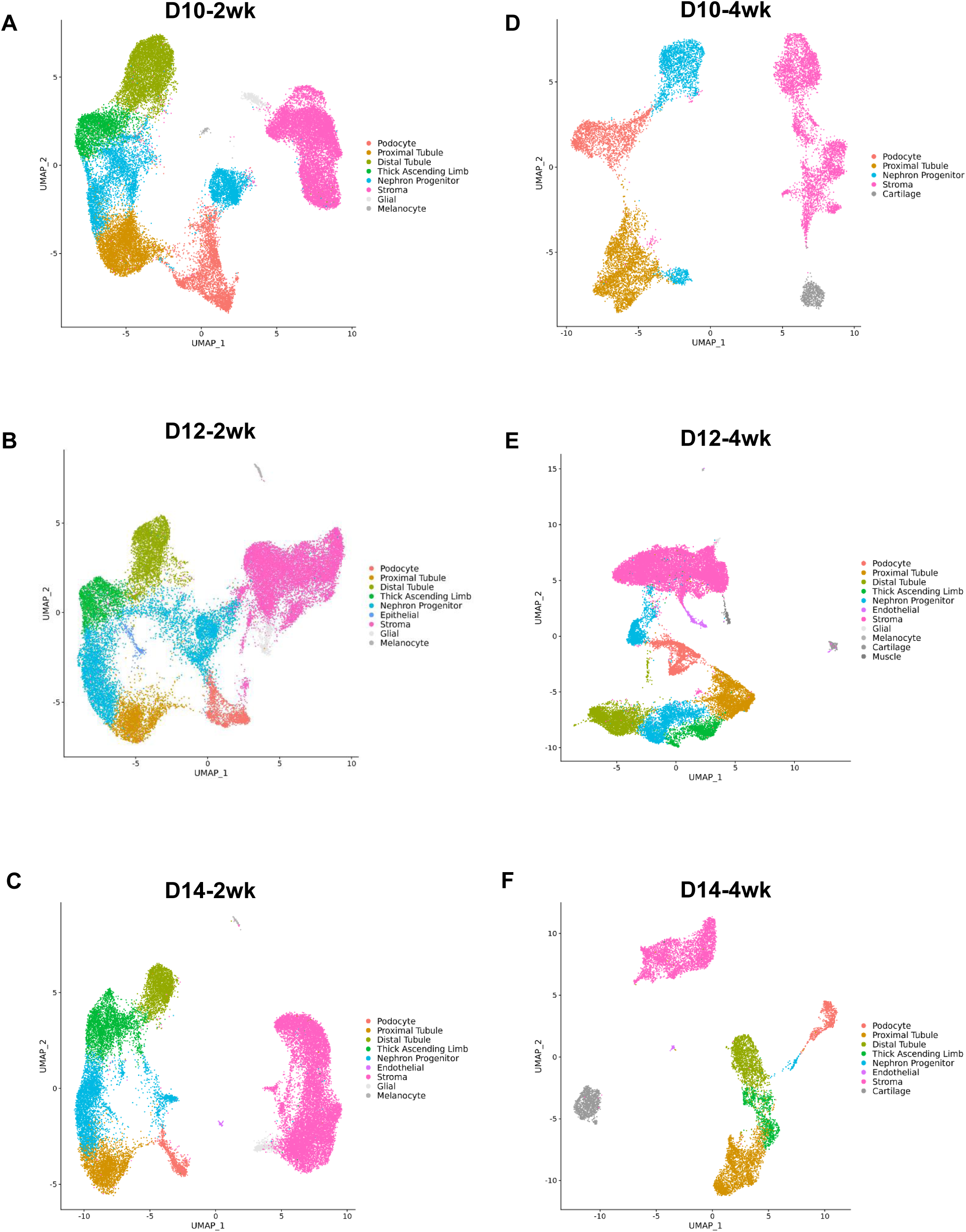
Two-dimensional representation of gene expression (Uniform Manifold Approximation and Projection, UMAP) of cell type populations for *in vivo* timepoints combined across replicates. (A) Day 10, 2-week maturation (D10-2wk) showing large populations of podocytes, tubular cells, and off-target cell types (melanocytes). (B) Day 12, 2-week maturation (D12-2wk) showing a small population of epithelial cells. (C) Day 14, 2-week maturation (D14-2wk) showing a small population of endothelial cells. (D) Day 10, 4-week maturation (D10-4wk) showing no populations of thick ascending limb or distal tubule cells, and a larger population of off-target cell types (cartilage). (E) Day 12, 4-week maturation (D12-4wk) showing more populations of off-target cell types (melanocytes, cartilage, muscle). (F) Day 14, 4-week maturation (D14-4wk) showing a large population of off-target cell types (cartilage).

**Supplementary Figure 4.**
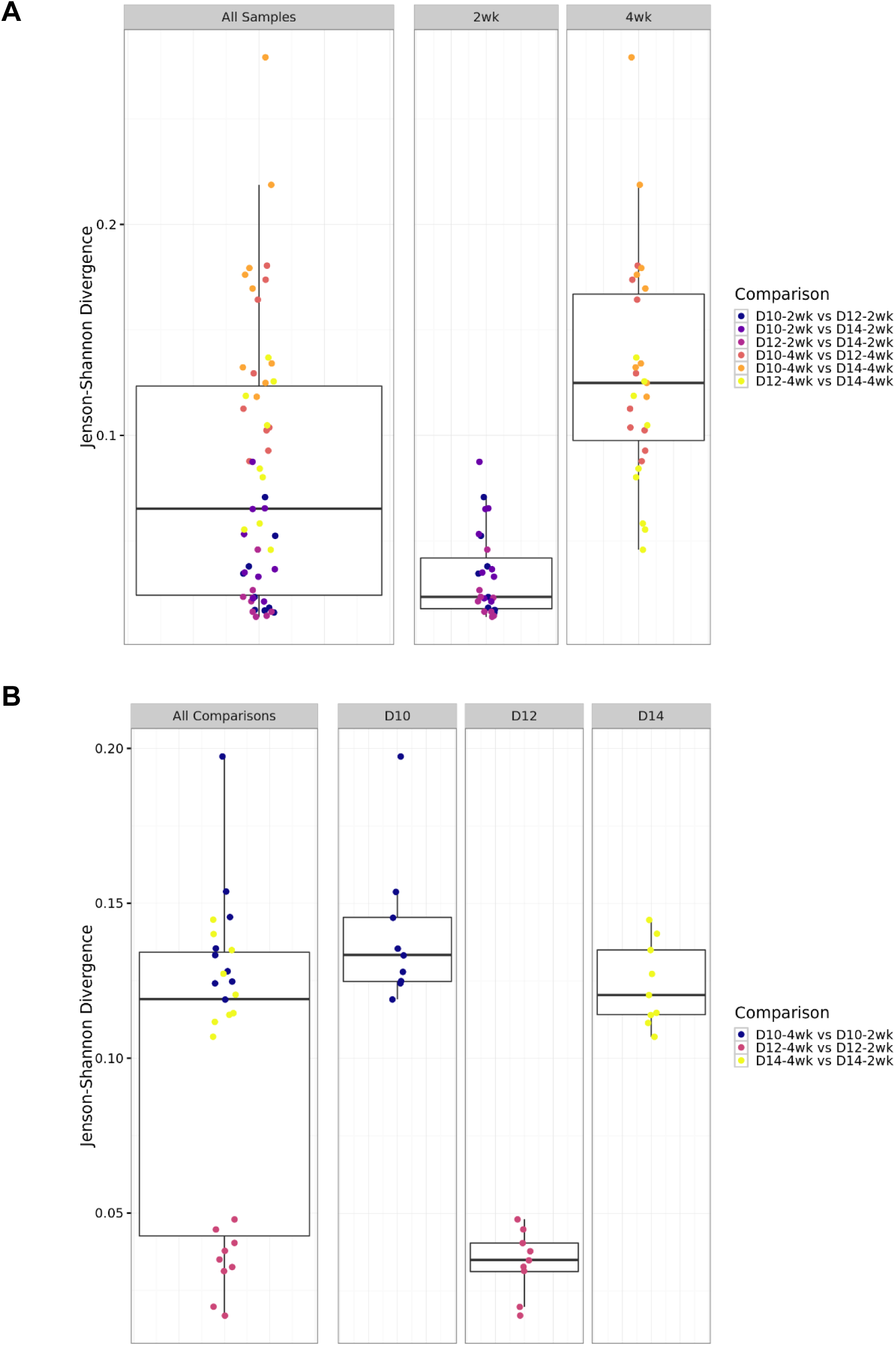
Comparison of cell type proportions for all *in vivo* timepoints using boxplots of Jenson-Shannon Divergence (JSD) indices. (A) Comparisons across transplantation days, by weeks of maturation reveals increased similarity after longer period of *in vivo* maturation. (B) Comparisons across weeks of maturation showing high similarity between weeks after transplantation at day 10 (D10) and day 14 (D14).

**Supplementary Figure 5.**
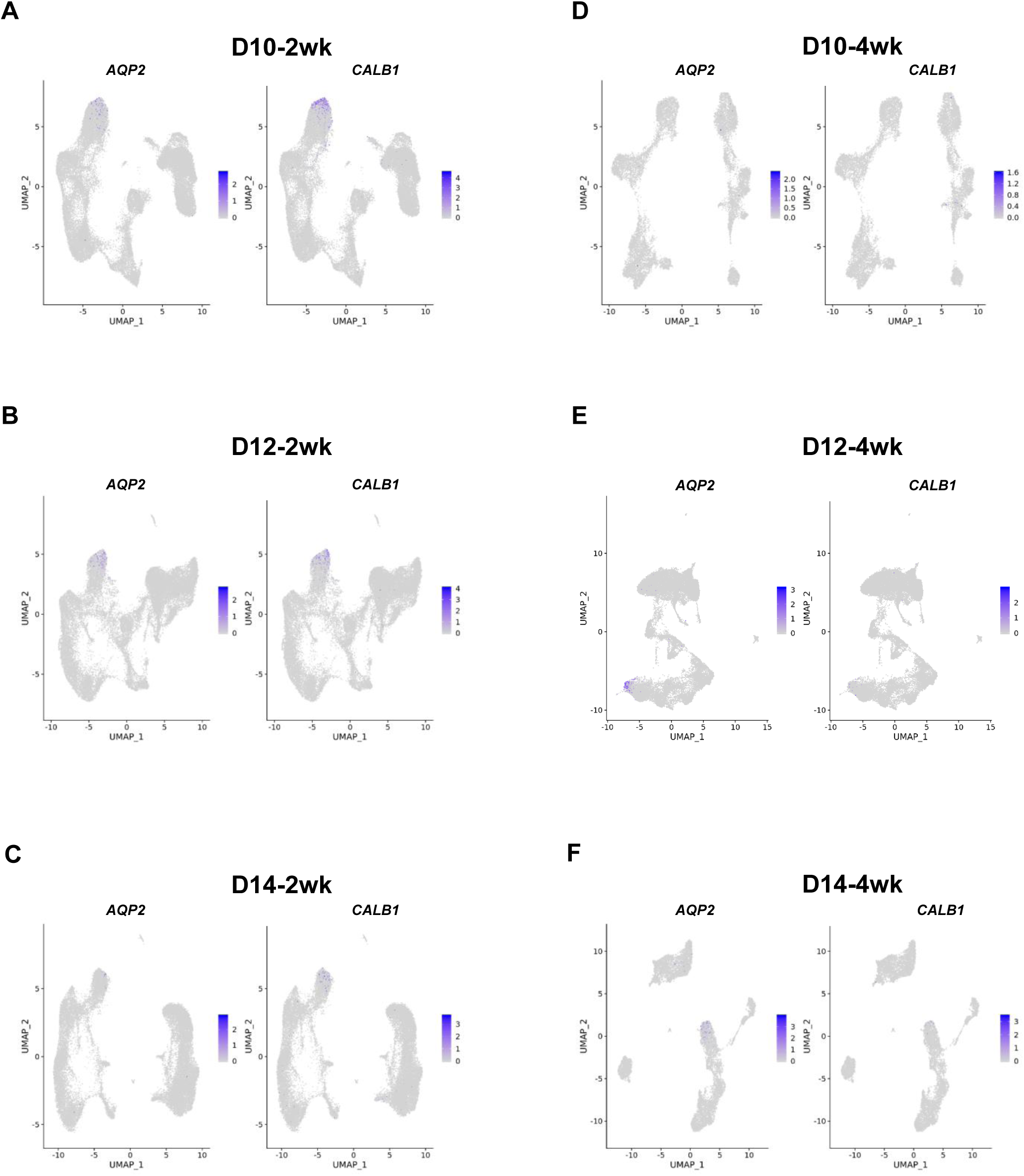
Two-dimensional representation of gene expression (Uniform Manifold Approximation and Projection, UMAP) of cell type populations for *in vivo* timepoints combined across replicates, showing cells expressing collecting duct marker genes *AQP2* (left) and *CALB1* (right) (A) D10, 2-week maturation (D10-2wk) organoids show highest population of cells expressing both markers. (B) D12, 2-week maturation (D12-2wk) organoids contain a subset of cells expressing both markers. (C) D14, 2-week maturation (D14-2wk) organoids show only a minority of cells expressing both markers. (D) D10, 4-week maturation (D10-4wk) organoids contain a minority of tubular cells expressing both markers. (E) D12, 4-week maturation (D12-4wk) organoids show a subset of tubular cells expressing both markers. (F) D14, 4-week maturation (D14-4wk) organoid contain a subset of cells expressing both markers.

**Supplementary Figure 6.**
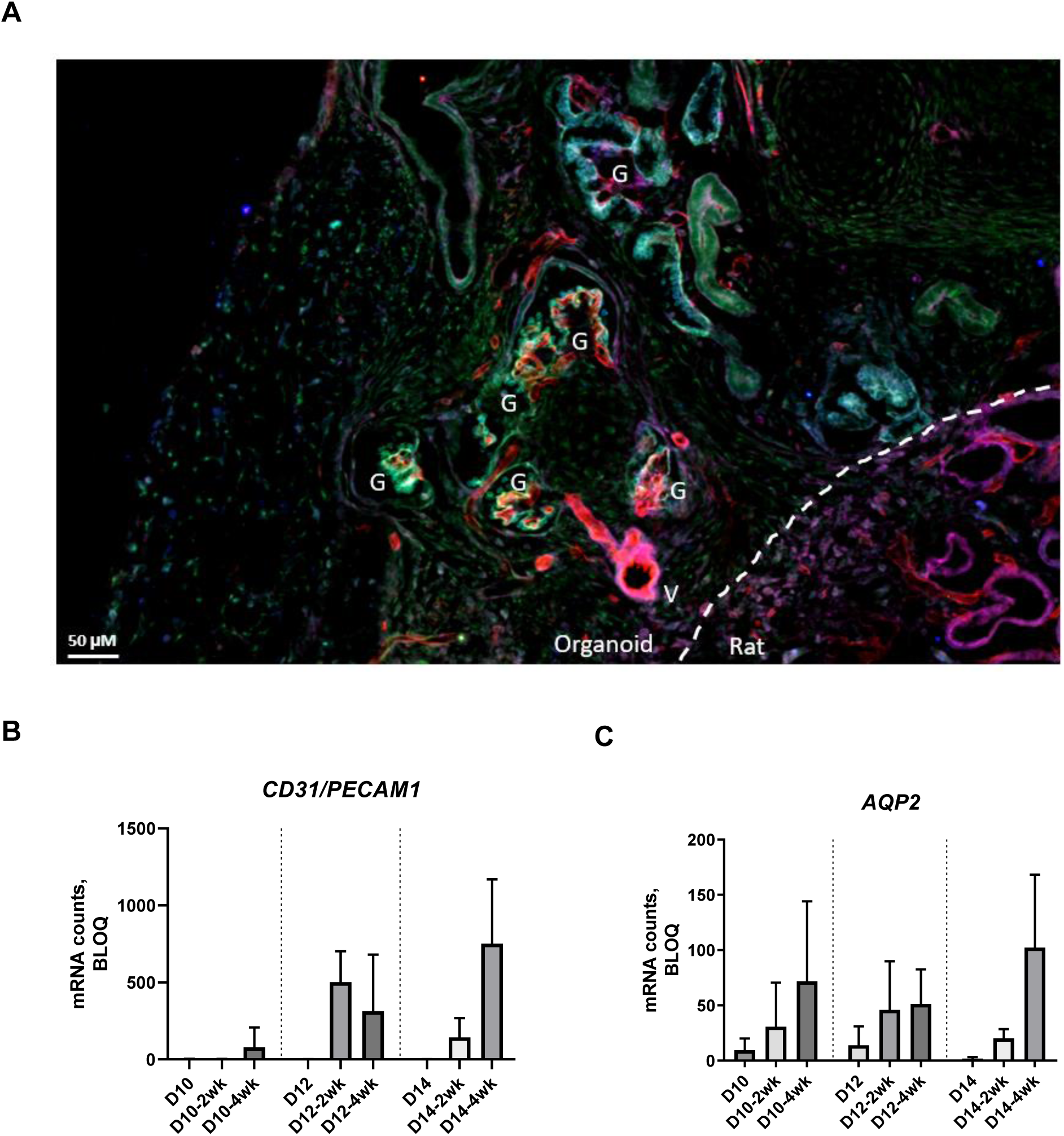
Transplantation promotes organoid vascularization and collecting duct maturation. (A) Representative transplanted organoid showing RECA-1 expressing rat derived vascularization of an organoid glomerulus. G: glomerulus; V: RECA-1 positive rat vessels. RECA-1: red; synaptopodin: green; nephrin: blue. (B) NanoString quantification of *CD31/PECAM1* induction over time for *in vitro* and transplanted organoids for d10, d12, and d14 plus 2wk or 4wk development *in vivo*. These data show that the there is significant endothelial *CD31/PECAM1* expression with transplantation. The most *CD31/PECAM1* expression occurs during D12 + 2-week, D12 + 4-week, and D14 + 4-week transplantation with no significant difference among these conditions (p>0.1). (C) NanoString quantification of *AQP2* induction over time for *in vitro* and transplanted organoids for D10, D12, and D14 plus 2-week or 4-week development *in vivo*. These data show that the there is significant endothelial A*QP2* expression with transplantation. The highest *AQP2* expression occurs during D10 + 2-week, D10 + 4-week, D12 + 2-week, D12 + 4-week, and D14+ 4-week transplantation.

**Supplementary Figure 7.**
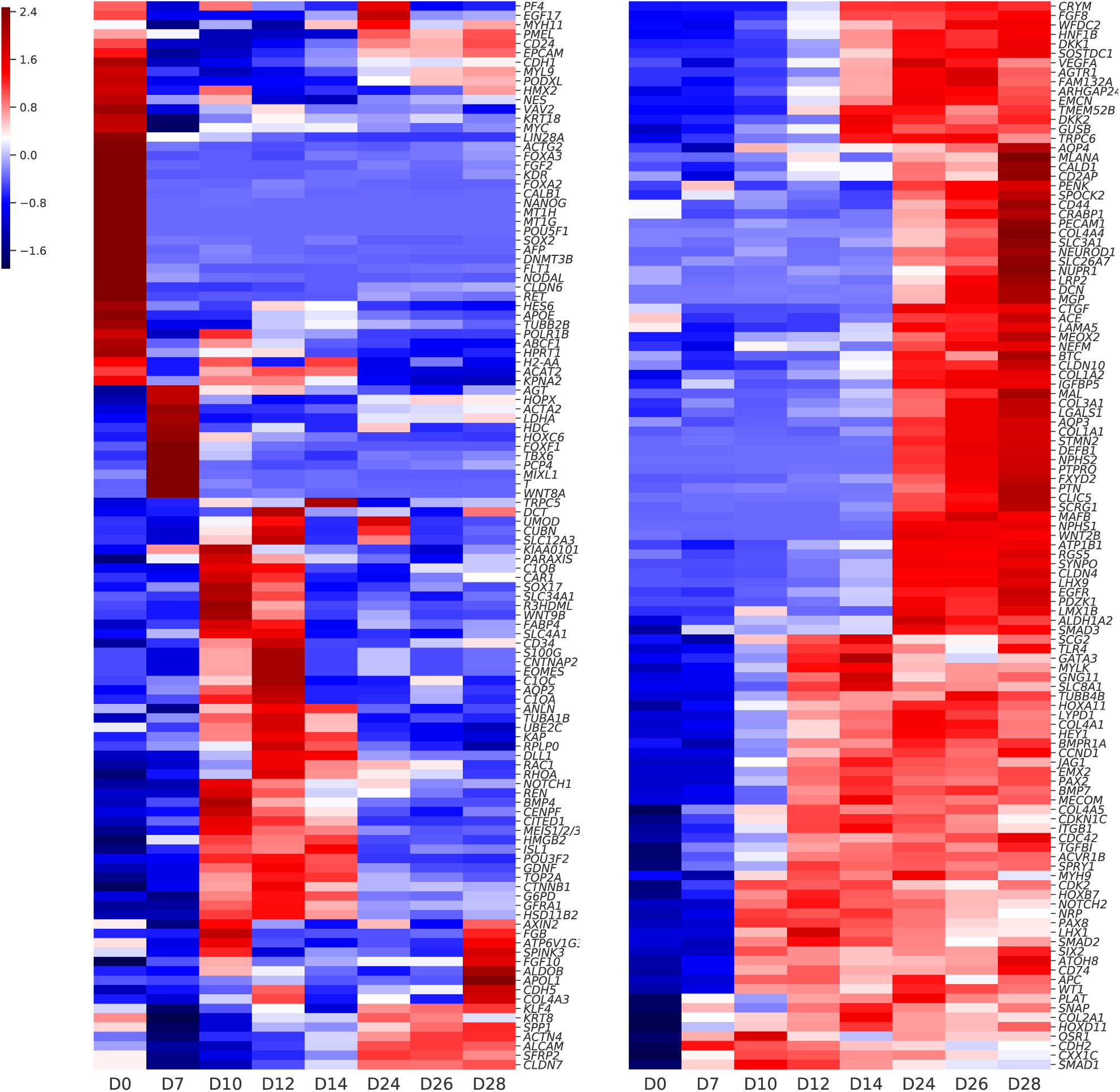
NanoString gene expression analysis. NanoString gene expression (row z-score) over *in vitro* organoid maturation (D0 through D28) for all 228 genes.

**Supplementary Figure 8.**
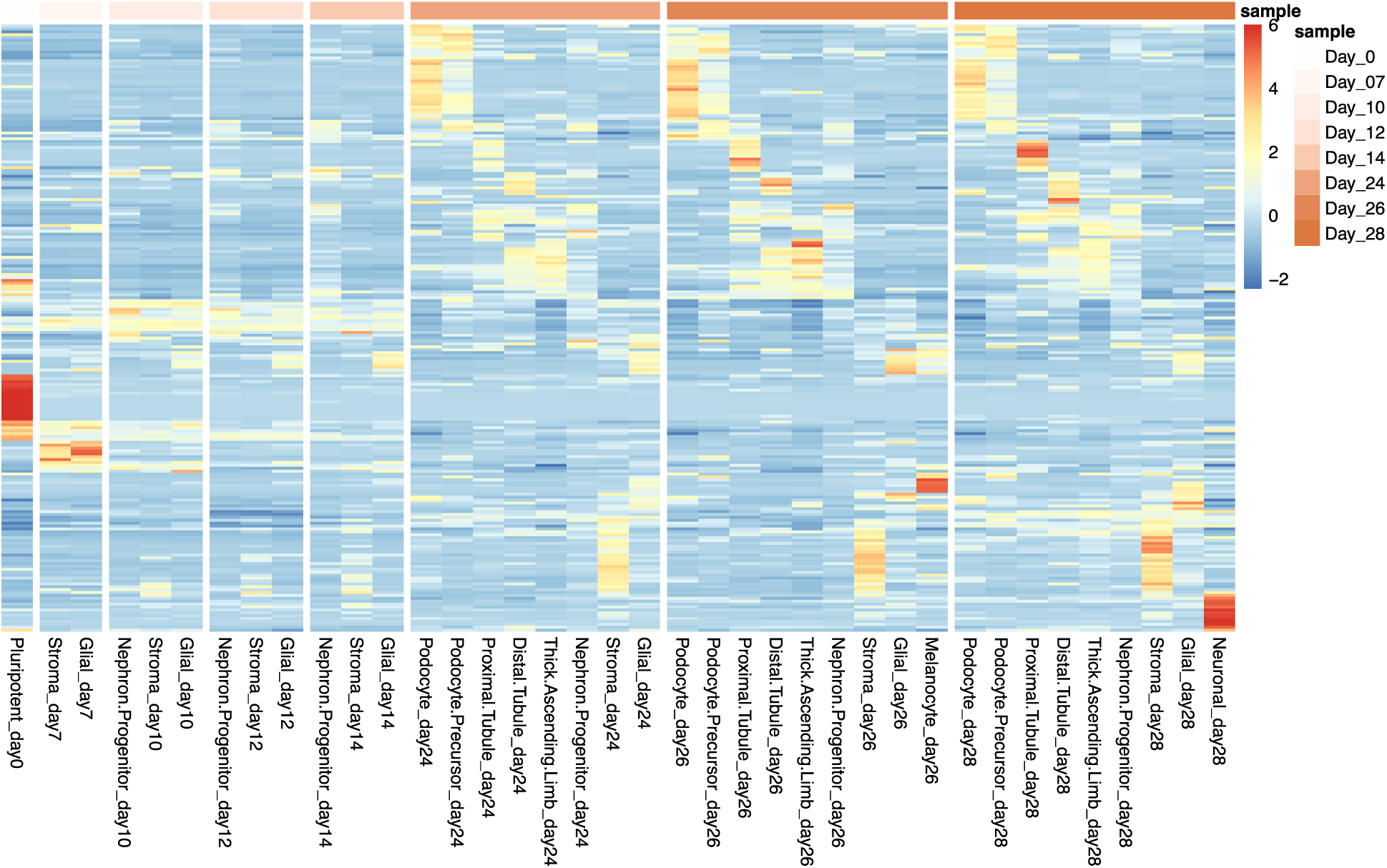
Expression analysis of NanoString marker genes in each cell type for all time points. Heatmap showing average gene expression scaled to row z-score within each scRNA-Seq cluster during *in vitro* organoid maturation (D0 through D28) for all 228 genes that were part of the NanoString panel.

**Supplementary Figure 9.**
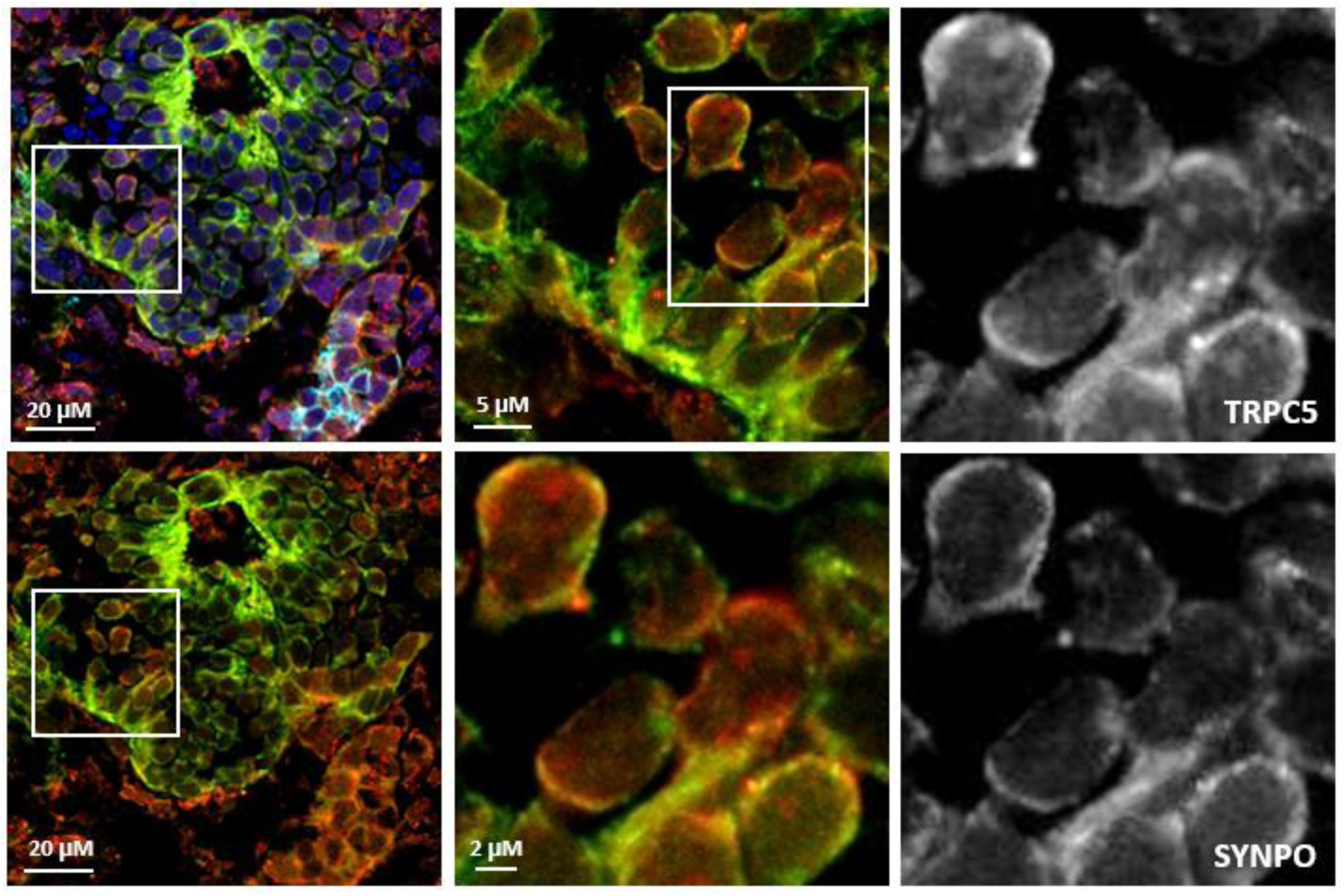
Double labeling with synaptopodin reveals podocyte TRPC5 protein expression in human organoids. Representative glomerulus (*left panel*) and zoomed in super-resolution images TRPC5 and synaptopodin staining in individual podocytes. Synaptopodin: green, TRPC5: red, Hoechst: blue.

Table 1 Description of 228 genes included in the iPSC and kidney organoid NanoString panel with annotations

